# Deficiency in POLE exonuclease causes synthetic lethality in highly aneuploid cancer cells

**DOI:** 10.1101/2025.07.16.665237

**Authors:** Eun Jung Kim, Alejandro Chibly, Kristel Dorighi, Jack Kamm, Szymon Juszkiewicz, Kevin Cabrera, Dexter Jin, Rohit Reja, Yangmeng Wang, Rachel Nakagawa, Minyi Shi, Yuxin Liang, Anwesha Dey, Ethan S. Sokol, Steven Gendreau, Meng He

## Abstract

Aneuploidy is a hallmark of cancer and is associated with drug resistance and poor clinical outcomes across diverse cancer types. However, no therapies have been clinically established to target highly aneuploid tumors. By analyzing a real-world dataset comprising nearly half a million tumor samples subjected to comprehensive genomic profiling, we identified a striking mutual exclusivity between POLE exonuclease domain mutations and high aneuploidy burden. This observation was independently validated using data from The Cancer Genome Atlas (TCGA) and the Cancer Cell Line Encyclopedia (CCLE). Probabilistic modeling revealed that the elevated quantity and unique spectrum of mutations induced by POLE exonuclease deficiency increase the likelihood of inactivating essential genes on chromosome arms harboring losses, leading to a synthetic lethal phenotype in highly aneuploid cells. Functional experiments demonstrated that POLE exonuclease activity is essential for the viability of highly aneuploid cancer cell lines but is dispensable in diploid cells. These findings suggest that selective inhibition of POLE exonuclease may represent a promising therapeutic strategy for targeting highly aneuploid tumors.

## Introduction

Aneuploidy is a cellular state characterized by the acquisition of an abnormal number of chromosomes or chromosome arms [1]. These chromosomal abnormalities are observed in over 90% of solid tumors and approximately 50% of hematological malignancies [2–4]. Aneuploidy has been recognized as both a consequence and a driver of tumorigenesis [5–7]. While aneuploidy often reduces fitness in normal cells, cancer cells tolerate and even exploit chromosomal imbalances for increased fitness [8–10]. Though the molecular mechanisms underpinning the initiation of aneuploidy is largely unknown, recent advances in single-cell genomics, karyotype-resolved sequencing, and functional genetics have illuminated the paradoxical role of aneuploidy in tumorigenesis [11]. Aneuploid genomes in cancer cells offer a substrate for clonal diversification under selective pressures, enabling tumor cells to adapt to microenvironmental stressors, evade immune detection, and resist therapeutic interventions [12,13]. Additionally, while chromosomal alterations can amplify oncogenes or eliminate tumor suppressors, the effects of chromosomal abnormalities can extend to global transcriptomic and proteomic disruption [14]. Previous work has implicated aneuploidy in the induction of proteotoxic and metabolic stress, disruption of mitotic fidelity, and activation of tumor-suppressive checkpoints, such as p53 [15–17].

Clinically, high aneuploidy burden, as well as specific recurrent aneuploidies—such as gain of chromosome 1q or 8q, or loss of 17p or 8p—are associated with poor prognosis, therapeutic resistance, and aggressive disease behavior across numerous cancer types, including breast, colorectal, ovarian, and lung cancers [10,18,19]. Mechanistically, aneuploidy promotes resistance through multiple routes, including increased intratumoral heterogeneity, reprogramming of apoptotic and stress response pathways, altered drug metabolism, and modulation of the tumor immune microenvironment [20]. For instance, tumors with high levels of aneuploidy often exhibit reduced antigen presentation and increased immunosuppressive signaling, features that correlate with diminished response to immunotherapies [21].

Therapeutic strategies targeting aneuploidy and consequential stresses have gained increased attention. Early preclinical efforts focused on amplifying the intrinsic vulnerabilities of aneuploid cells, such as proteotoxic and mitotic stress, through the use of proteasome inhibitors, heat shock protein (HSP) inhibitors, or spindle assembly checkpoint modulators [22–24]. More recently, synthetic lethality screens have identified aneuploidy-specific dependencies, including elevated reliance on autophagy, DNA repair pathways, and the unfolded protein response [25,26]. For example, cells with whole-chromosome gains are more sensitive to centrosome clustering inhibitors or agents targeting oxidative phosphorylation and energy homeostasis [27]. Despite promising preclinical results, clinical translation remains challenging. Many stress pathway inhibitors lack tumor specificity or induce systemic toxicity [28]. Nonetheless, early-phase trials have evaluated compounds such as the HSP90 inhibitor ganetespib and proteasome inhibitor bortezomib, with mixed results [22,23,29].

In this study, we analyzed a real-world database comprising half a million patient samples across over 400 cancer types and subtypes to identify potential synthetic lethal targets for high aneuploidy burden. We identified and validated the exonuclease domain of POLE as a potential therapeutic target for tumors exhibiting high aneuploidy burden.

## Results

### Characterization of chromosomal arm-level aneuploidy in a real-world comprehensive genomic profiling database

We examined a real-world database of comprehensive genomic profiling results derived from 499,320 patient samples. This database encompasses over 90 different primary cancer types, including non-small cell lung cancer, colorectal cancer, breast cancer, pancreatic cancer, ovary cancer, prostate cancer, etc. (Figure 1A). Patient and tumor characteristics, including chromosomal sex, histological cancer type, local/metastasis biopsy site and genetic ancestry, are well represented in this database (Figure 1B).

**Figure 1:**
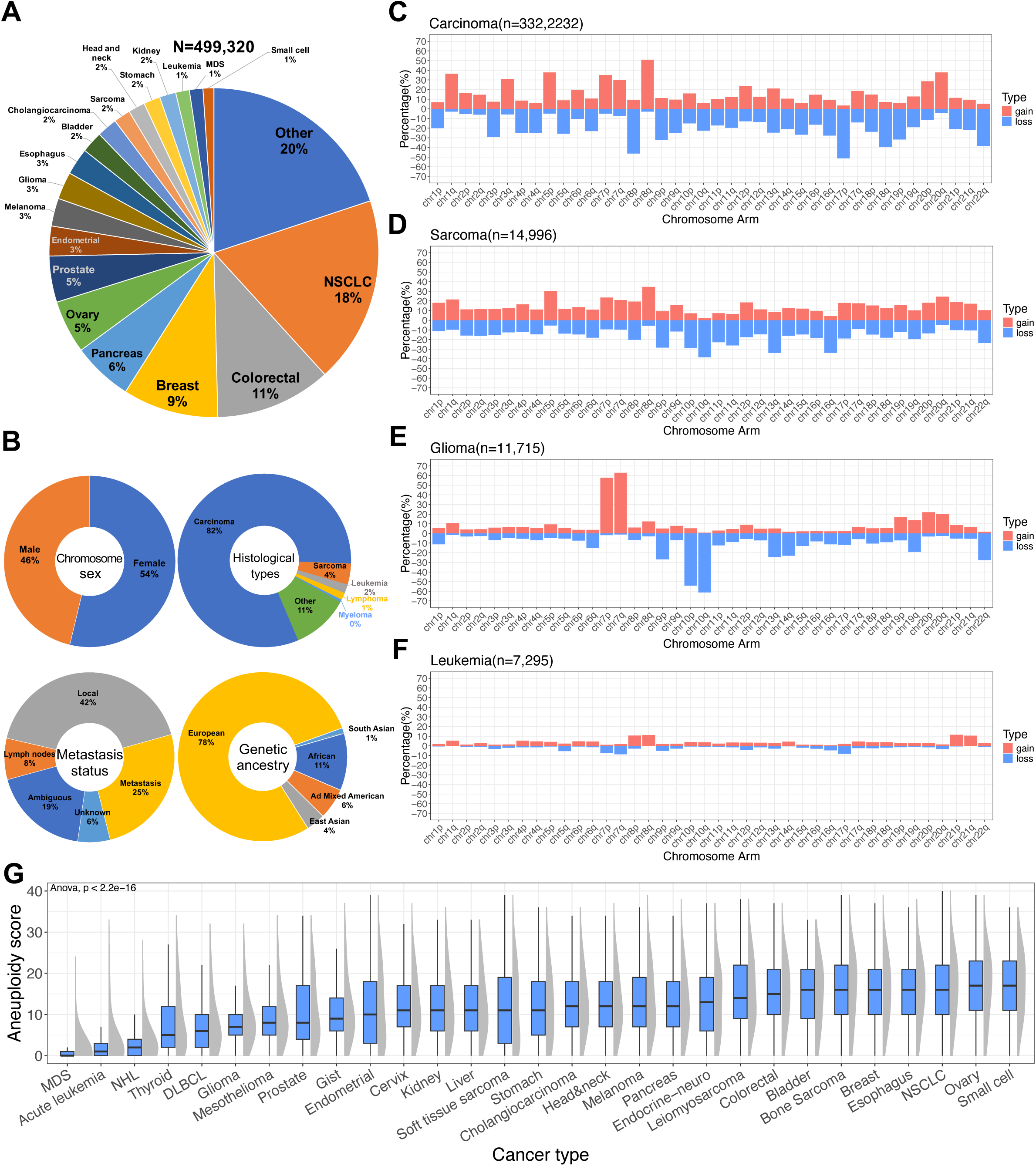
Characterization of chromosomal arm level aneuploidy in the real-world comprehensive genomic profiling database. (A) Pie chart summarizing the real-world database by primary cancer type. (B) Donut charts summarizing the real-world database by chromosome sex, histological types, metastasis status and genetic ancestry. (C-F) Frequencies of each chromosomal arm loss or gain in carcinoma (C), sarcoma (D), glioma (E) and leukemia (F). (G) Distribution of aneuploidy scores across major cancer types (In each boxplot, the center line represents the median value. The lower and upper hinges correspond to the first and third quartiles, respectively, while the whiskers extend up to 1.5x the interquartile range. This applies to all the boxplots presented in this study).

A computational algorithm was run to call chromosomal arm level gains and losses for 40 autosomal chromosomal arms for each sample [30,31]. The arm-level aneuploidy caller revealed unique chromosome alterations across different histological cancer types (Figure 1C-1F). Consistent with previous reports, 8p loss, 17p loss, 1q gain and 8q gain are particularly prevalent in carcinomas, as compared to sarcomas, gliomas and leukemias [3,18,19], whereas whole chromosome 7 gain and whole chromosome 10 loss were observed as unique features in gliomas [32]. On the other hand, leukemia exhibited significantly less arm-level aneuploidy [33]. Upon examining the arm-level aneuploidy across different primary tumor types (Supplemental Figure 1), we also observed distinct distributions of specific chromosomal alterations. For instance, while 1q gain is common across various carcinoma cancer types, it is particularly common in breast cancer with a near 70% incidence rate. 3p loss is particularly enriched in small cell, head and neck, cholangiocarcinoma and kidney cancers with an incidence rate over 50%. Another notable example is 13q gain which is detected in approximately 70% of colorectal cancer but is rare in most other cancer types.

To estimate the aneuploidy burden in each tumor type, we calculated an aneuploidy score by counting the total number of chromosome arm losses or gains in a tumor sample [3,34,35]. Not surprisingly, we observed pronounced differences in aneuploidy scores across tumor types (Figure 1G). Hematological malignancies such as leukemia, non-Hodgkin lymphoma, and diffuse large B-cell lymphoma exhibit significantly less aneuploidy. Even within solid tumor types, gliomas, mesotheliomas, and prostate cancers tend to have significantly lower aneuploidy scores compared to breast cancer, non-small cell lung cancer and small cell cancer.

Additionally, we observed statistically significant differences in aneuploidy scores among patients with different genetic ancestries (Supplemental Figure 2A). Patients of Admixed American ancestry generally display lower aneuploidy compared to patients of African ancestry. Higher aneuploidy burden has been associated with metastasis [36,37], which was consistent with our observation of significantly higher aneuploidy scores in tumor samples collected from metastatic sites (Supplemental Figure 2B). A strong correlation between aneuploidy scores and patient age was also noted (Supplemental Figure 2C). Chromosomal arm losses and gains contribute equally to aneuploidy scores, and no correlation was observed between these two types of alterations (Supplemental Figure 2D), suggesting these events are likely selected independently during cancer evolution.

### Elastic net regression model identifies genetic mutation of POLE as a top negative predictor of aneuploidy score

To systematically evaluate the impact of patient and tumor characteristics, including sex, tumor subtype, genetic ancestry, metastasis status, and genomic alterations on aneuploidy, we deployed an elastic net regression model using aneuploidy score as the dependent variable (see Method for details). The comprehensive results from the elastic net regression model can be found in Supplemental Table S1, with an optimal alpha value of 0.25 (Supplemental Figure 3A). Genomic alterations with non-zero coefficients are summarized in Figure 2A. The model identified *TP53* alteration as the top positive predictor of aneuploidy score, a well-established driver of cancer genomic instability and aneuploidy [4,6,38]. In non-transformed primary cells, *TP53* inactivation is often required for cells to tolerate aneuploidy [8,39,40]. Thus, *TP53* alteration serves as a positive control, demonstrating the robustness of our model. Further examination showed that tumors with homozygous loss (CN) of *TP53* exhibit significantly higher aneuploidy scores than those with *TP53* short variants (SV) or structural rearrangements (RE) (Figure 2B and Supplemental Figure 3B). Apart from *TP53*, our model identified several other genomic alterations as top positive predictors for aneuploidy scores, such as *AR, GID4, AKT3, and FGF10*. Although the functional link between these genomic alterations and aneuploidy remains unclear, it is interesting to note that alterations of *GID4, AKT3,* and *FGF10* are predominantly through amplification (Supplemental Figure 3C), and the association between *AR* and aneuploidy score is driven by *AR* amplification in prostate cancer (Supplemental Figure 3D).

**Figure 2:**
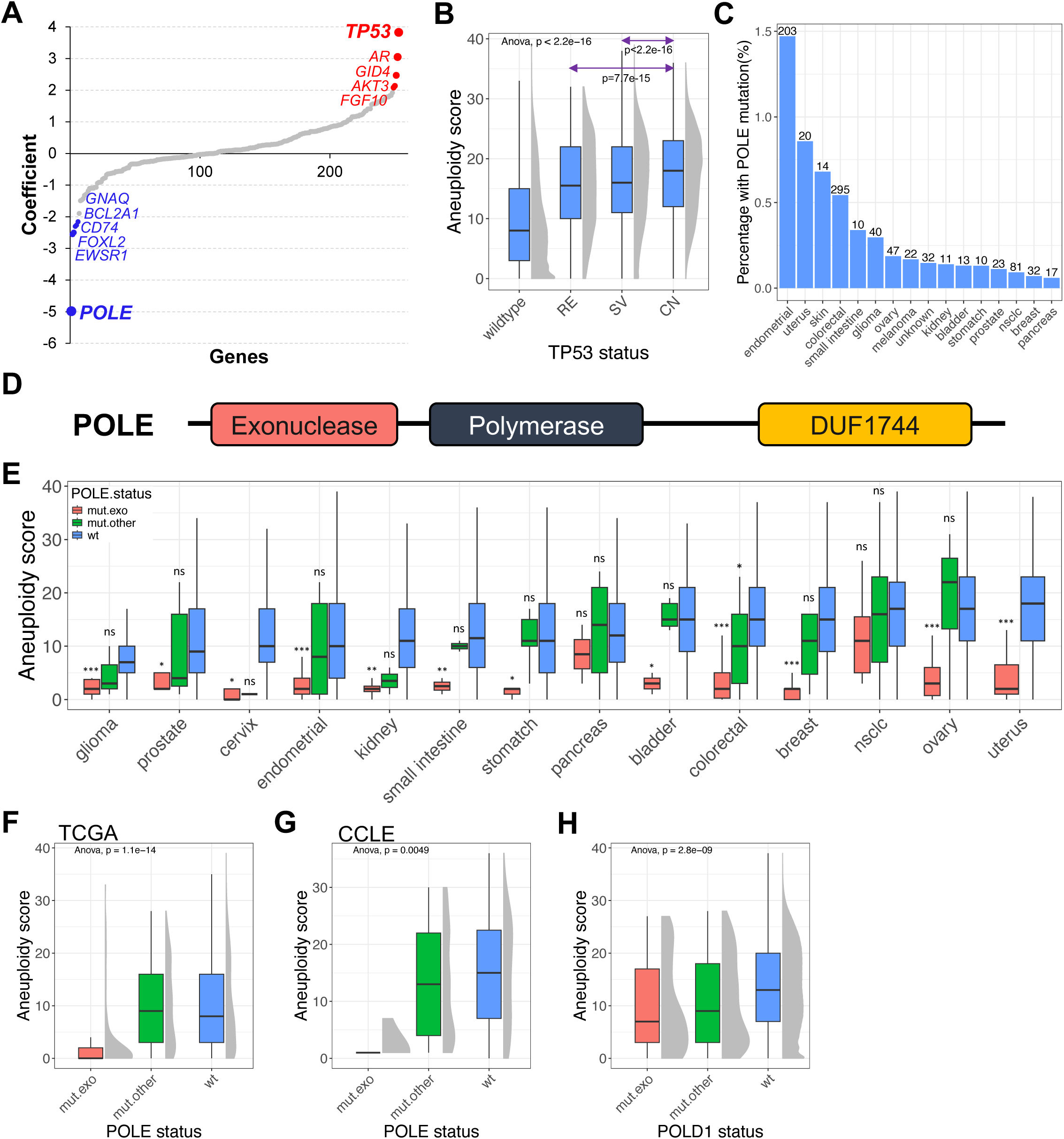
Mutations in POLE exonuclease domain are mutually exclusive with high aneuploidy burden. (A) Elastic net regression model identified genes with non-zero coefficient to be predictors for aneuploidy scores. Top positive predictor genes are highlighted in red, while top negative predictor genes are highlighted in blue. (B) Distribution of aneuploidy scores across tumors with different *TP53* alterations. RE, structural rearrangement; SV, short variant; CN, copy number alteration (homozygous deletion). Student t-tests were used for two group comparisons. (C) Frequency of *POLE* mutation across various cancer types in the real-world database. (D) Representation of POLE protein domains. (E) Distribution of aneuploidy scores in tumors with different *POLE* alterations across various cancer types in the real-world database. mut.exo, mutations within the POLE exonuclease domain (a.a.268-a.a.471); mut.other, mutations outside the POLE exonuclease domain; wt, wild type. Student t-tests were used for two group comparisons (mut.exo vs. wt or mut.other vs. wt), * p<0.05, ** p<0.01, *** p<0.001, ns p>0.05. (F) Distribution of aneuploidy scores in tumors with different *POLE* alterations in TCGA. (G) Distribution of aneuploidy scores in cell lines with different *POLE* alterations in CCLE. (H) Distribution of aneuploidy scores in tumors with different *PODL1* alterations in the real-world database.

The model also identified several negative predictors for aneuploidy scores, such as *GNAQ, BCL2A1, CD74, FOXL2, EWSR1*, and notably *POLE*. Although their functional links with aneuploidy are unclear, the strong signal that alterations in *POLE* is associated with lower aneuploidy scores warranted further investigation. *POLE* encodes the catalytic subunit of the DNA polymerase epsilon complex and is most frequently mutated in endometrial (1.5%) and colorectal cancers (0.5%) (Figure 2C). *POLE* encodes a protein with multiple functional domains, including the exonuclease domain, the DNA polymerase domain, and the less characterized DUF1744 domain [41,42] (Figure 2D). Hotspot mutations in the *POLE* exonuclease domain have been reported [41,43] and separating tumors with mutations within this domain versus outside of the domain revealed that the negative association with high aneuploidy burden is driven predominantly by mutations within the exonuclease domain (Figure 2E and Supplemental Figure 3E). This association was validated in The Cancer Genome Atlas (TCGA) (Figure 2F) and the Cancer Cell Line Encyclopedia (CCLE) (Figure 2G). Across all three datasets, tumors with exonuclease domain mutations of *POLE* are near diploid compared to those with mutations outside the exonuclease domain or *POLE* wild-type tumors.

We also looked to see if there was an association between aneuploidy score and *POLD1* alterations. *POLD1* encodes the catalytic subunit of DNA polymerase delta. Both *POLE* and *POLD1* are essential B-family DNA polymerases in eukaryotic cells, playing crucial roles in DNA replication and repair [44–47]. Both enzymes possess 5′ to 3′ polymerase and 3′ to 5′ exonuclease proofreading activities, ensuring high-fidelity DNA synthesis. However, they differ in their primary functions during replication: POLE is primarily responsible for leading strand synthesis, while POLD1 mainly synthesizes the lagging strand [44]. Mutations in the exonuclease domains of POLE and POLD1 can compromise their proofreading abilities, leading to increased mutation rates [41,42]. Interestingly, our elastic net regression model identified *POLE* alteration as a top negative predictor to aneuploidy scores, but not *POLD1*. We therefore examined the association between aneuploidy scores and *POLD1* mutation status. While there was a statistically significant reduction in aneuploidy score in *POLD1* exonuclease (a.a.304-a.a.533) mutated tumors, the effect size was minimal and less consistent across cancer types compared to *POLE* (Figure 2H and Supplemental Figure 3F).

### Deficiency in POLE exonuclease causes synthetic lethality in highly-aneuploid cells by inactivating essential genes on chromosome arms with losses

We were interested in understanding the underlying mechanism that drives the mutual exclusivity between POLE exonuclease mutations and high aneuploidy burden in tumor cells. Mutual exclusivity in biology has been proposed to stem from either functional redundancy or synthetic lethality [48]. POLE exonuclease mutations are often associated with ultra mutated tumors, characterized by extremely high tumor mutation burdens (TMB), in many cases exceeding 500,000 mutations per haploid genome (Figure 3A and Supplemental Figure 4A) [45]. Although there is no clear association between TMB and aneuploidy scores when examined across all tumors (Supplemental Figure 4C) [3], we observed a strong negative correlation between TMB and aneuploidy scores in *POLE*-mutated tumors (Figure 3B). POLE exonuclease mutations are not only associated with ultrahigh mutations but also associated with a unique mutational signature [49,50], leading us to the hypothesis that the ultrahigh TMB and unique mutation spectrum caused by POLE exonuclease mutations lead to synthetic lethality in tumor cells with high aneuploidy burden (Figure 3C). We proposed that in highly aneuploid tumor cells with numerous chromosome arm losses and gains, the inactivation of POLE’s exonuclease domain would lead to ultrahigh mutations across the genome. The unique spectrum associated with these mutations may have a high probability of inactivating essential genes located on chromosome arms with losses, leading to cancer cell death (Figure 3C).

**Figure 3:**
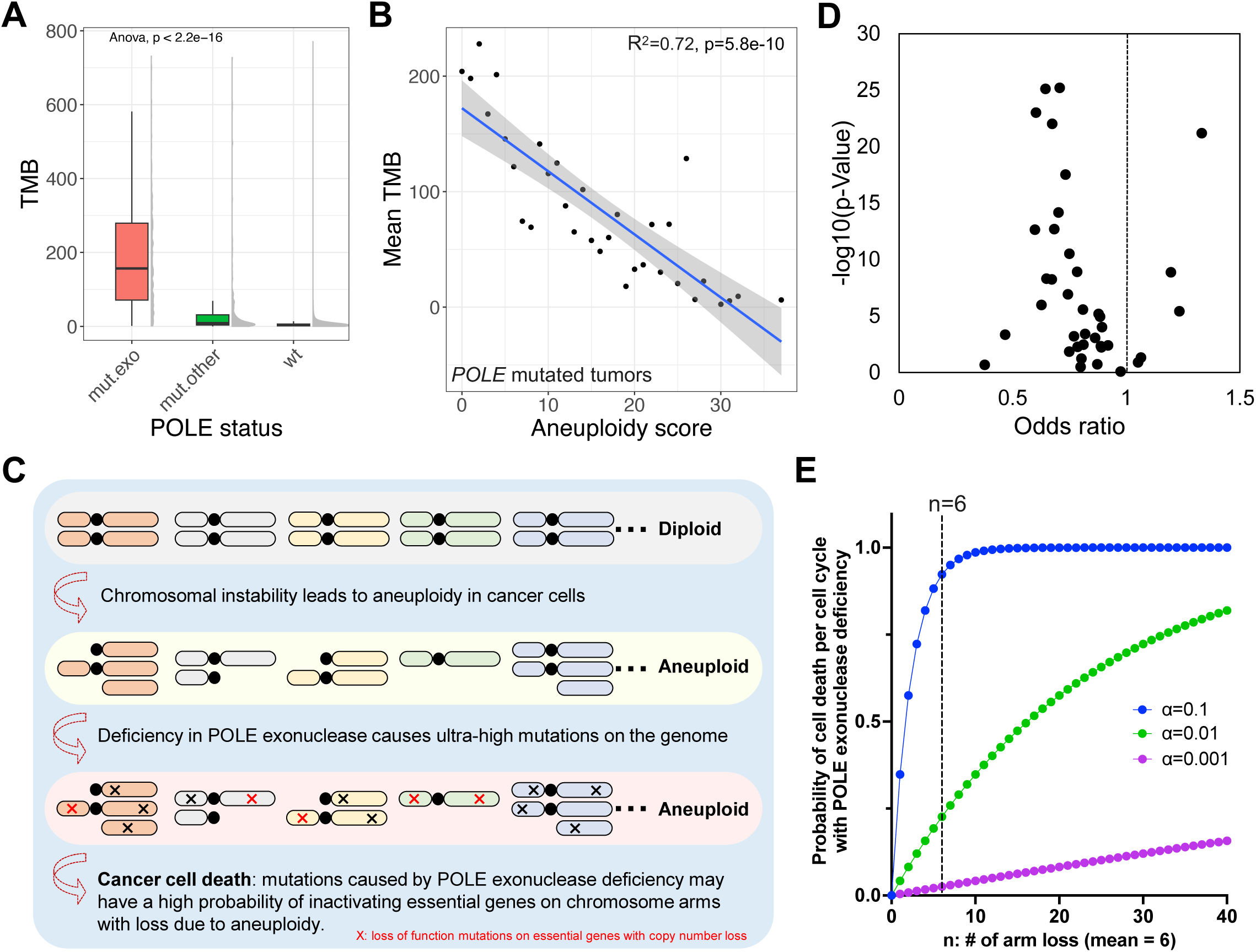
High mutation burden and the unique mutation spectrum caused by POLE exonuclease deficiency have a high probability of inactivating essential genes in highly aneuploid cells. (A) Distribution of tumor mutational burden (TMB) in tumors with different *POLE* alterations in the real-world database. (B) Aneuploidy scores and the mean TMB are negatively correlated in *POLE*-mutated tumors (blue line: a linear regression model fit by lm() function in R) (C) Illustration of proposed mechanism of mutual exclusivity between high aneuploidy burden and POLE exonuclease mutations: mutations caused by POLE exonuclease deficiency have a high probability of inactivating essential genes on chromosome arm harboring losses in highly aneuploid cells. (D) Most of the chromosomal arms are less likely to have co-occurrence of arm losses and genetic alterations (limited to variants with unknown significance), odds ratio < 1. Each dot represents an autosomal chromosomal arm. (E) Probabilistic modeling of cell death per cell cycle in cells with POLE exonuclease deficiency. *Pr_cell death per cell cycle_ ≈ 1 – exp (-n × P_arm_ × α × m)*, *n =* number of chromosome arm losses (mean = 6), *P_arm_ =* probability of a mutation (caused by POLE exonuclease deficiency) hitting an essential gene divided by total number of chromosome arms (*P_arm_=0.00057*), *α =* percentage of POLE exonuclease deficiency-induced mutations on essential genes that are deleterious (unknown), *m* = number of mutations caused by POLE exonuclease deficiency per cell cycle per haploid genome (*m=7500* estimated for stage IV colorectal cancer), refer to methods section for more details.

Consistently, in POLD1 exonuclease mutated tumors where we did not observe strong mutual exclusivity with high aneuploidy burden (Figure 2H), the TMB is substantially lower in comparison to POLE exonuclease mutated tumors (Supplemental Figure 4B). Consistent with our hypothesis, we observed in the real-world database that chromosomal arm losses and random mutations (limited to variants with unknown significance) are less likely to co-occur (odds ratio < 1) (Figure 3D). Next, we developed a probabilistic model to estimate the likelihood of mutations caused by POLE exonuclease deficiency causing lethality by inactivating essential genes (see Method for details).

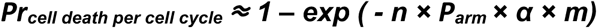

In this model, **n** stands for the number of chromosome arms with losses in a tumor cell, ***P_arm_*** stands for the average probability of a mutation caused by POLE exonuclease deficiency hitting an essential gene per chromosome arm, **α** stands for the portion of deleterious mutations hitting essential genes, and it remains a challenge to determine the true value of **α**. Lastly, **m** stands for the number of mutations POLE exonuclease deficiency can cause in a cell per cell cycle per haploid genome. Based on this model, we determined that for a typical solid tumor cell with an average of 6 chromosome arm losses (based on the real-world database), if 10% (**α**) of mutations caused by POLE exonuclease deficiency hitting an essential gene are deleterious, the probability of inactivating at least one essential gene, leading to cell death per cell cycle, approaches 100%. If we assume **α** = 0.01, the probability of cell death per cell cycle under the same condition is approximately 25% (Figure 3E). As the value of **α** becomes smaller, the probability of cell death per cell cycle is less likely to induce a synthetic lethality or mutual exclusivity phenotype.

The elastic net regression model also identified microsatellite instability as another top negative predictor for aneuploidy score (Supplemental Table 1, Supplemental Figure 4F). Mutational signatures related to mismatch repair deficiency also negatively correlate with low aneuploidy burden (Supplemental Figure 4D). In previous reports, a synthetic lethal interaction between microsatellite instability and high aneuploidy burden was also proposed, however, the underlying mechanisms are unknown [51–53]. Interestingly, even though microsatellite unstable tumors exhibit low aneuploidy burden, almost comparable to POLE exonuclease mutated tumors (Supplemental figure 4F), these tumors showed significantly lower TMB compared to POLE exonuclease-mutated tumors (Supplemental Figures 4E and 4G), but more comparable to POLD1 exonuclease-mutated tumors. However, the TMB in POLD1 exonuclease-mutated tumors are likely not sufficient to drive the mutual exclusivity with high aneuploidy burden (Figure 2H). Therefore, it remains to be determined if the mechanism we proposed here for POLE can be applied to microsatellite unstable tumors.

### POLE exonuclease activity is essential for highly-aneuploid cancer cells but not diploid cells

While our probabilistic modeling provided evidence that the mutual exclusivity between POLE exonuclease mutations and high aneuploidy burden in tumor cells are likely driven by synthetic lethality, the unknown value of the key variable **α** undermines the model’s validity. Next, in order to experimentally validate that POLE exonuclease activity is required for survival of highly-aneuploid cancer cells, but not diploid cells, we utilized an inducible adenine base editor to perform a mutagenesis screen using guide RNAs targeting POLE’s exonuclease domain (Figure 4A) [54]. We selected four cancer cell lines with high aneuploidy scores (EFE184, MDST8, SW480, SW837) and four diploid cell lines (DLD1, HCT116, HEC265, LS180) (Figure 4B). We first validated the inducible expression of the adenine base editor across a few cell lines, including a non-cancerous cell line 293, an aneuploidy cell line SW837 and a diploid cell line DLD1(Supplemental Figure 5A). Then, we performed a mutagenesis screen on the eight cancer cell lines (Figure 4C, see Method for details). Seven of the eight cell lines propagated post-lentiviral transduction, except for MDST8 (Figure 4D). Expression of the adenine base editor was maintained during the screen (Figure 4E), and guide RNAs were amplified at passage 8, sequenced, and quantified. Guide RNAs targeting POLE’s exonuclease domain were consistently underrepresented in the three highly aneuploid cell lines (EFE184, SW480, SW837), but not in the diploid cell lines (DLD1, HCT116, HEC265, LS180) (Figure 4F, Supplemental Figure S5). This supports our hypothesis that POLE exonuclease domain inactivation induces synthetic lethality in highly aneuploid cells but not in diploid cells. Consistent with the base editor screen, we showed that, in the highly aneuploid cell line SW480, ectopic expression of wild type POLE rescued the fitness defect caused by *POLE* knockdown, while the two hot-spot mutations in the POLE exonuclease domain (P286R and V411L) failed to rescue the defect (Figure 4G, H). Together, our experimental data across multiple cancer cell line models demonstrated the essentiality of POLE exonuclease activity in highly aneuploid cells.

**Figure 4:**
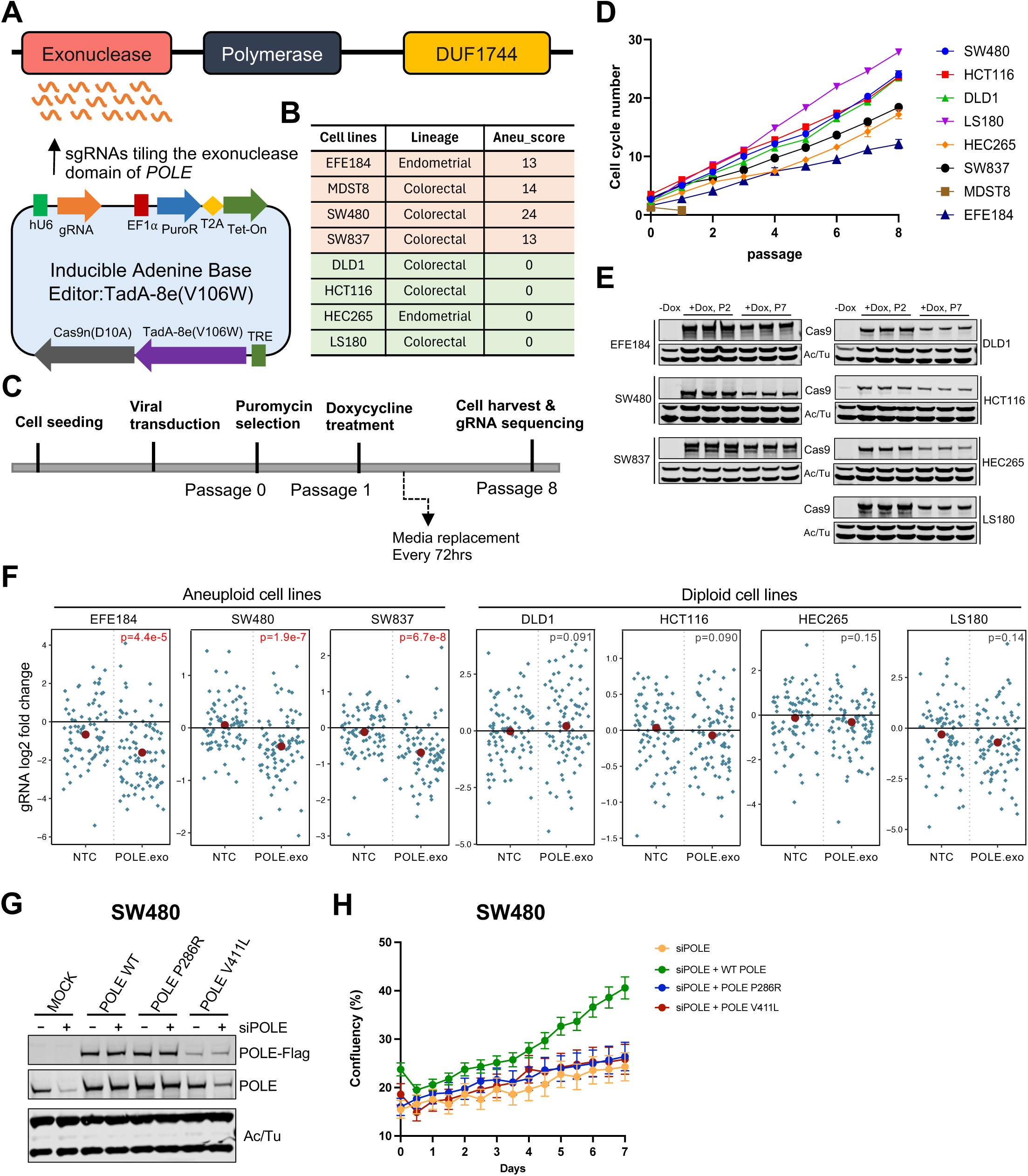
Mutagenesis screening validated that POLE exonuclease is essential for the survival of highly aneuploid cells but not diploid cells. (A) Illustration of the inducible adenine base editor screen tiling the exonuclease domain of POLE. (B) Cell lines used for the adenine base editor screen. (C) Experimental flow of the adenine base editor screen. (D) Cell growth curves post viral transduction and induction of adenine base editor. (E) Western blot confirming the sustained expression of the adenine base editor system during the experiment. (F) Compared to non-targeting control gRNAs (NTC), gRNAs tiling the exonuclease domain of POLE are significantly underrepresented in the aneuploid cell lines (EFE184, SW480, SW837), but not in the diploidy cell lines (DLD1, HCT116, HEC265, LS180). P values and log2 fold change were determined by comparing guide RNA counts at passage 8 to passage 0. (G) Western blot confirming the siRNA mediated knockdown of endogenous POLE, and over expression of ectopic POLE variants (WT, P286R and V411L) in SW480 cell line. (H) Growth curves of SW480 cells under different genetic backgrounds.

## Discussion

Developing therapies for cancers with high aneuploidy burden is of high medical importance. Synthetic lethality is an attractive strategy for identifying therapeutic targets, due to the unknown molecular mechanisms causing aneuploidy and the complex biological aberrations associated with it. In this study, we leveraged a real-world cancer genomics database and identified that mutations in the POLE exonuclease domain are mutually exclusive with high aneuploidy burden. We hypothesized and validated that POLE exonuclease inactivation causes synthetic lethality in highly aneuploid cells, likely by inactivating essential genes with copy number losses. This discovery leads to a therapeutic hypothesis that selective inhibition of POLE exonuclease may benefit patients with highly-aneuploid tumors.

Previous reports indicate that endometrial or colorectal cancer patients with POLE-mutated tumors often have favorable clinical outcomes, even without immune checkpoint inhibitors [55,56]. The mechanisms underlying this prognosis are not fully understood. One hypothesis suggests that these tumors may exhibit high tumor antigens, enhancing immune system recognition [57]. We speculate that the near-diploidy nature of *POLE*-mutated tumors could also contribute to these favorable clinical outcomes. Additionally, selective inhibition of POLE exonuclease might potentiate immune checkpoint inhibitors by enhancing tumor antigens through hypermutation.

However, inhibiting POLE’s exonuclease domain carries risks. This domain plays a critical role in genome duplication proofreading for both cancer and normal cells. Individuals with germline mutations in POLE’s exonuclease domain often develop Polymerase Proofreading-Associated Polyposis (PPAP) and have a higher risk of colorectal and uterine cancers later in life [45,46]. Interestingly, these patients do not exhibit strong acute disease symptoms. Murine models show that heterozygous inactivation of POLE exonuclease presents no discernible phenotypes compared to wild-type animals, while homozygous inactivation correlates with higher cancer incidence, similar to patients with heterozygous mutations [58]. The hotspot somatic mutation P286R exhibits a pronounced tumorigenesis phenotype in animal models that is not observed with germline mutations [59]. This aligns with yeast studies where analogous substitutions induced strong mutator phenotypes exceeding those of conventional proofreading mutations [60]. The introduction of a charged residue within the exonuclease site appears to sterically prevent channeling of the nascent DNA strand containing an erroneous base towards the exonuclease active site residues [61], instead likely facilitating continuous DNA synthesis (Supplemental Figure 6A). Similarly, V411L substitution lies within the ‘wedge helix’ directly involved in the separation of erroneous base-containing nascent DNA strand from the template during the transfer from polymerase to exonuclease site (Supplemental Figure 6B) [62]. A longer side chain of leucine compared to valine is likely to push forward the wedge helix, creating a stronger physical barrier preventing efficient transfer of the leading strand from the polymerase to the exonuclease site. This might explain the high prevalence and hypermutator phenotype of the V411L mutation in cancers.

Although there is precedent for developing small molecule inhibitors of RNase H-like exonucleases [63], it remains to be determined whether similar pharmacological interventions could induce hypermutator activity in POLE and what the therapeutic implications of such an approach might be. Despite the safety and druggability challenges associated with POLE exonuclease, our work proposes a novel concept of introducing synthetic lethality in highly aneuploid tumor cells. Aneuploidy might expose many essential genes as vulnerabilities due to large-scale chromosome losses and gains. Strategies to inhibit these essential genes might offer therapeutic opportunities.

## Methods

### Key resources table

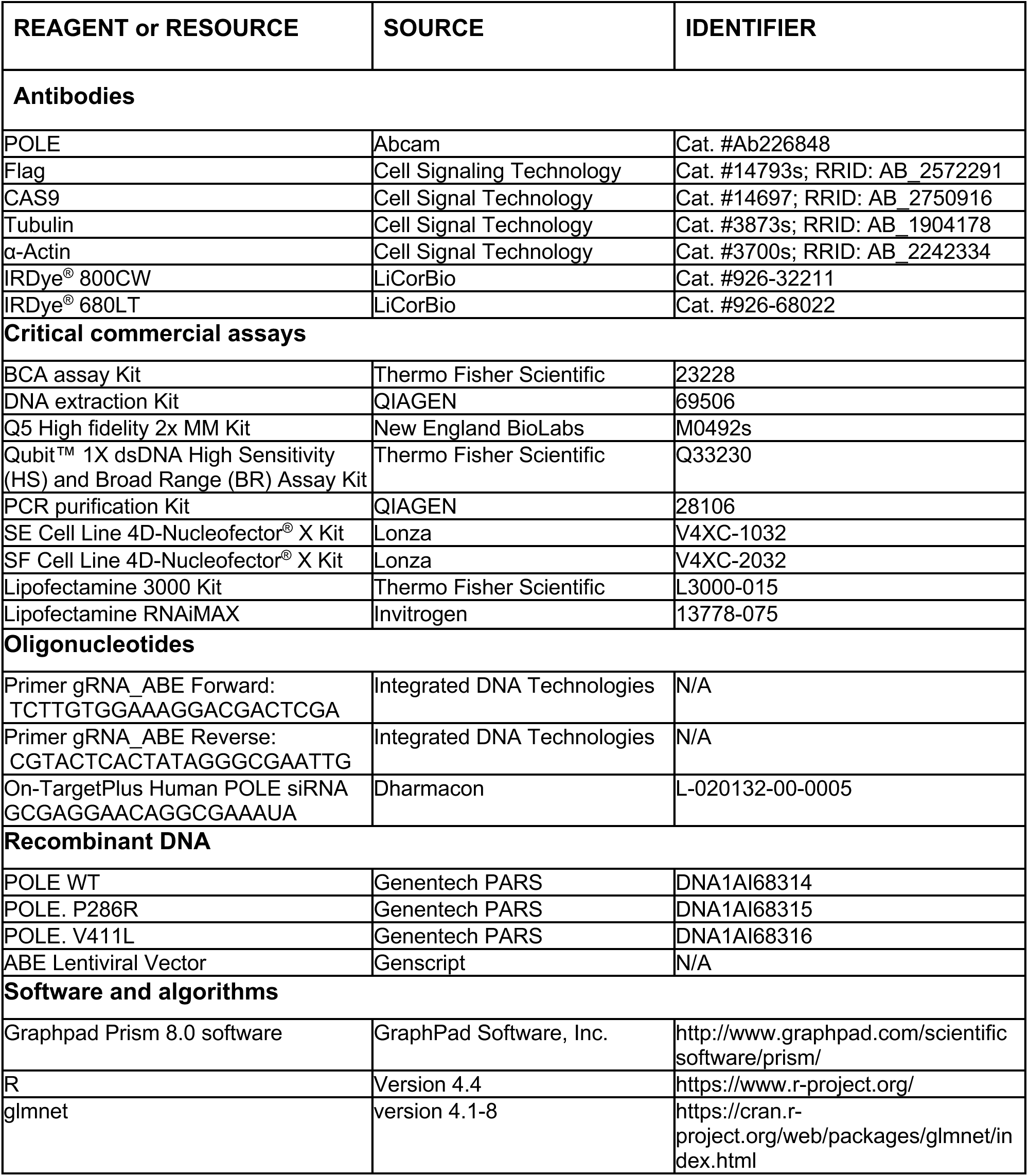

### Comprehensive genomic profiling in a real-world database

Comprehensive genomic profiling of formalin-fixed, paraffin-embedded (FFPE) tissue sections of 499,320 tumor samples during routine clinical care was performed using the FDA-approved FoundationOne®CDx or the laboratory developed tests, FoundationOne® or FoundationOne®Heme in a Clinical Laboratory Improvement Amendments–certified, College of American Pathologists–accredited laboratory (Foundation Medicine Inc.). Hybrid capture was carried out on at least 300 cancer-related genes and select introns from 28-34 genes frequently rearranged in cancer. Arm level gains and losses were called in a research use only capacity based on statistically lower or higher log2 CN for at least 50% of the arm, as previously described [31]. Approval for this study, including a waiver of informed consent and Health Insurance Portability and Accountability Act (HIPAA) waiver of authorization, was obtained from the Western Institutional Review Board (protocol no. 20152817).

### Elastic net regression

The R package glmnet version 4.1-8 (https://cran.r-project.org/web/packages/glmnet/index.html) was used to fit an elastic net regression model using aneuploidy score (0-40) as a dependent variable. The following features were selected as predictor variables: disease ontology terms (n=487), metastasis status (n=2), chromosome sex (n=2), genetic ancestry (n=5), microsatellite instability (n=2) and genetic alterations of evaluable genes (n=349). The outputs of the predictor variables were converted to binary (0,1). The real-world dataset was split into a training set and a test set with a ratio of 2:1. To identify the optimal model, a total number of 21 different alpha values (0-1) were tested with an interval of 0.05. Mean Squared Error (MSE) was calculated for each alpha value to determine the performance of each model. Alpha value with the smallest MSE was chosen for the final model.

### Probabilistic modeling of process of inactivating essential genes

Deficiency in POLE exonuclease leads to a unique mutational signature [49,50], with approximately 30% of the mutations on trinucleotide motif TCT and 20% of the mutations on TCG. This unique mutation spectrum allows us to estimate the probability of a mutation caused by POLE exonuclease deficiency hitting an essential gene. The list of essential genes in mammalian cells were extracted from a previous study [64]. First, we compute the probability of a mutation caused by POLE exonuclease deficiency falling on the genomic coding region for essential genes on each autosomal chromosome arm, through each trinucleotide motif:

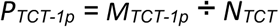

*N_TCT_* = total number of the trinucleotide motif TCT on human genome

*M_TCT-1p_* = total number of the trinucleotide motif TCT on coding regions for all essential genes located on chr1p

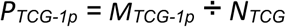

*N_TCG_* = total number of the trinucleotide motif TCG on human genome

*M_TCG-1p_* = total number of the trinucleotide motif TCG on coding regions for all essential genes located on chr1p

Similar calculations can be done for all trinucleotide motifs.

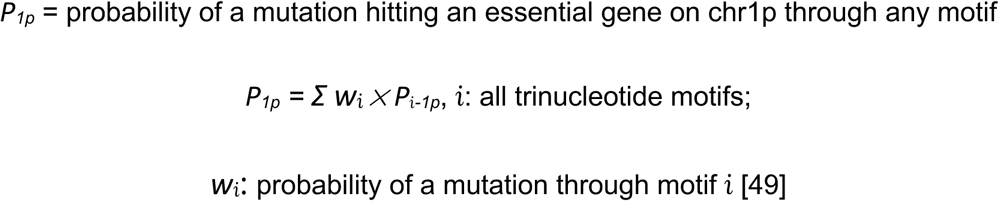

TCT and TCG are the dominant motifs [49], we therefore can simplify the equation above

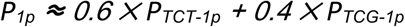

Similarly, the probabilities for all the 40 autosomal chromosome arms can be calculated as:

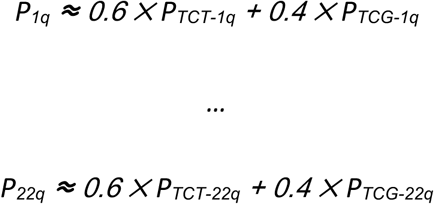

The values of P*_1p_* - P*_22q_* can be computed from the trinucleotide count table in supplementary table 2. In order to evaluate the probability as a function of chromosome arm loss and other variables, we computed an average P*_arm_* that will be used for all chromosomal arms in diploid cells.

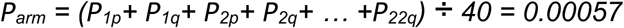

Even though we can compute the probability of a mutation hitting essential genes for each chromosome arm, it remains unknown what portion of those mutations, when hitting an essential gene, will lead to loss of function. We therefore introduced a variable ***α***, proportion of mutations on essential genes that are deleterious, to control the impact.

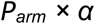

Lastly, we need to determine the number of mutations (**m**) tumor cells can accumulate per cell cycle per haploid genome when POLE exonuclease is deficient. This number was estimated using the same approach as previously reported [65]. Briefly, the number of cell cycles (**cc**) required for a single POLE exonuclease mutated cancer cell to give rise to a tumor were calculated based on the average tumor volume for stage IV colorectal cancer (45cm^3^) [66], and the average volume of a mammalian cell (1.8*×*10^-6^ mm^3^) [65]. The variable **m** is then calculated as

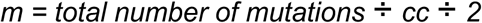

The average total number of mutations in POLE exonuclease mutated tumors is approximately 540,000 (TMB *×* 3,000) per genome based on the real-world data and TCGA, and the number of cell cycles (**cc**) is approximately 36, based on which we computed **m** = 7500. However, the TMB is estimated from a bulk of cancer cells, this estimation assumes that tumor cells are not evolutionally divergent, therefore this could be an overestimation of **m**. In a POLE exonuclease deficient tumor cell with **n** chromosome arm losses, the probability of inactivating at least one essential gene with copy number loss per cell cycle is:

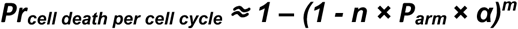

As ***m*** is large and ***n × P_arm_ × α*** is small, this equation can be simplified as

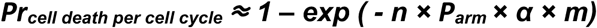

which follows from the Poisson approximation to the Binomial distribution, or equivalently the fact that (1 − *p*)*^m^* → exp (−*λ*) when *m* → ∞, *pm* → *λ*. This formula could alternatively be derived by assuming that mutations occur as a Poisson process with total rate **m** per cell cycle, of which ***n × P_arm_ × α*** are lethal.

We note that the approximation above contains several simplifications that affect its accuracy. First, there is a wide variance in the observed TMB across POLE-exonuclease deficient samples, and thus in **m**. Secondly, different samples with the same **n** may have different numbers of haploid bases depending on which chromosome arms are lost. Therefore, it would improve accuracy to account for the specific TMB and aneuploidy pattern for any given sample. Additionally, both **m** and **n** are estimated from bulk rather than single-cell data. Different cells within a sample may have different values of **n**, causing overdispersion; meanwhile, the estimated TMB includes mutations that are not present in all cells, which may inflate **m**, since it is based on the number of mutations per cell divided by the number of cell cycles.

### Western blots

Whole cell lysates were extracted using RIPA buffer (50 mM Tris HCl, 150 mM NaCl, 1.0% (v/v) IGEPAL CA-630, 0.5% (w/v) Sodium Deoxycholate, 1.0 mM EDTA, 0.1% (w/v) SDS and 0.01% (w/v) sodium azide at a pH of 7.4) supplemented with protease and phosphatase inhibitors (Thermo Fisher Scientific, 78427,1861279). Protein concentrations were determined using the BCA assay. Equal amounts of protein (20 µg per lane) were separated on 4-12% acrylamide gels (Invitrogen, wg1403bx10) and transferred to membranes (Bio-Rad 1704159) using a transfer system (trans-blot turbo, Bio-Rad). Membranes were blocked in 5% BSA (Gibco,15260-037) in TBST (Tris-buffered saline with 0.1% Tween-20) for 30 minutes at room temperature, followed by an overnight incubation at 4°C with primary antibodies diluted in blocking buffer at 1:500. After washing three times with TBST, membranes were incubated with secondary antibodies for 1 hour at room temperature. Protein bands were visualized using Odyssey CLx Infrared Imaging System.

### Inducible adenine base editor screen

Tad-8e(V106W) was cloned downstream of a TRE3G doxycycline-inducible promoter in a lentiviral expression vector with the EF1a promoter driving a puromycin resistance gene linked to Tet-ON 3G. The vector also contains a human U6 promoter driving a single guide RNA cassette into which the guide RNA library containing 89 POLE exonuclease tiling guide RNAs, 85 non-targeting control guide RNAs and 46 putative positive control guide RNAs were cloned. All cloning was performed by Genscript. To perform the screen, 10 million cells were seeded in 15 cm plates and infected with the adenine base editor lentiviral library at a multiplicity of infection (MOI) of 0.2. After 48 hours, the medium was replaced with a fresh growth medium containing 3 μg/ml puromycin for 3–5 days. Following antibiotic selection, cells were trypsinized and re-seeded at a density of 2 million cells per 10 cm dish in triplicates, with doxycycline added to induce base editor expression. Cells were maintained for 8 passages, cell pellets containing 2 million cells were collected at the end of the experiment and before doxycycline treatment (as baseline).

### DNA Isolation and gRNA amplification

Genomic DNA was isolated from 2 million cells collected from the adenine base editor screen using the QIAGEN DNA extraction kit, following the manufacturer’s protocol. gRNAs were amplified by PCR using the C1000 Touch Thermocycler (Bio-Rad). The PCR amplification was performed using a standard three-step thermocycling protocol for 25 cycles: denaturation at 95 °C, primer annealing at 55 °C, and extension at 72 °C, employing the Q5 High-Fidelity 2X Master Mix (New England BioLabs, M0492S). The resulting amplicons were purified using a PCR purification kit (Qiagen, 28106) for downstream sequencing.

### gRNA amplicon library preparation and sequencing

For generation of gRNA amplicon libraries, the KAPA HyperPrep Kit (Roche) that has incorporated custom adapters and library amplification PCR primers from Integrated DNA Technologies was used with an input of 40ng of amplicon DNA. Amplicon libraries were quantified, and the average library size was determined using the D1000 ScreenTape on 4200 TapeStation (Agilent Technologies). Libraries were pooled and the Qubit dsDNA HS Assay Kit (Thermo Fisher Scientific) was used to quantify the pool. Library pools were sequenced on NextSeq 2000 (Illumina) to generate 3 million paired-end 51-base pair reads for each sample.

### Analyses of base editor screen

Gene-level significance for differential barcode abundance was assessed using the FRY test implemented in the limma R package (https://bioconductor.org/packages/release/bioc/html/limma.html). Each cell line was evaluated individually comparing the gRNA abundance targeting POLE exonuclease versus a set of non-targeting control (NTC) gRNAs. For each gene, multiple gRNAs targeting that gene were considered as a predefined set, and the FRY method was used to estimate a log2-Fold change in barcode abundance with a corresponding p value.

### POLE knockdown and rescue experiments

To establish cell lines stably expressing different *POLE* variants (WT, P286R, and V411L) in SW480 cells, we transfected the cells with plasmids encoding human *POLE* variants using electroporation. Briefly, 10^5 cells were suspended with 2 μg plasmids in the electroporation buffer (SE and SF cell line 4D X kits, Lonza Bioscience). Electroporation was carried out using optimized settings per the manufacturer’s protocol (Lonza, 4D Nucleofector X unit). Post-transfection, cells were plated in complete growth medium for recovery. Stable expression of POLE was verified by western blot following puromycin selection for three weeks. For rescue experiments, endogenous POLE expression was silenced using siRNA (5’-GCGAGGAACAGGCGAAAUA). Cells were transfected with a siRNA mixture containing 20 nM POLE-targeting siRNA and RNAiMAX transfection reagent (Invitrogen, Cat# 13778-075) according to the manufacturer’s protocol. Twenty-four hours post-transfection, 2,000 cells were seeded into a 96-well plate and monitored by the Incucyte system (Essen BioScience) every 12 hours over a period of seven days. Incucyte’s built-in software tools were employed to evaluate cell confluence.

### Resource availability

#### Lead contact

Further information or requests for data and resources should be directed to, and will be fulfilled by the lead contact, Dr. Meng He

#### Materials availability

The all-in-one inducible adenine base editor system generated for this study can be requested through the standard Genentech MTA process. Please reach out to the lead contact for details.

#### Data and code availability

Public datasets used in this study include TCGA and CCLE datasets, which are available for download from cBioPortal (https://www.cbioportal.org/) and DepMap (https://depmap.org/portal/), respectively. The sequencing data for the real-world database are derived from clinical samples. All the relevant data supporting the findings of this study are provided within the main manuscript and its supplementary files. Due to HIPAA requirements, we are not authorized to share individualized patient genomic data, which contains potentially identifying or sensitive patient information. Foundation Medicine is committed to collaborative data analysis, with well-established, and widely utilized mechanisms by which investigators can query our core genomic database of >700,000 de-identified sequenced cancers to obtain aggregated datasets. For more information and mechanisms of access, please contact the corresponding author(s) or the Foundation Medicine, Inc. Data Governance Council at. All computational codes used in the analysis are available for research purposes and can be provided upon request.

## Acknowledgments

This study is funded by F. Hoffmann-La Roche AG

## Author contributions

M.H. and S.G. contributed to the conception of the study. E.S., K.C., D.J. and S.M. contributed to the curation of the real-world database. M.H. performed analysis. E.J.K. performed the wet lab experiments. A.C. analyzed the base editor screen. K.D. designed the base editor system and contributed to the interpretation of the data. S.J. performed the structural analysis of POLE. R.R. and J. K. contributed to the statistical modeling. Y.W. and R. N. contributed to the interpretation of the data and the writing and revision of the manuscript. M.S. and Y.L. generated library from the gRNA library and performed next generation sequencing. A.D. contributed to the interpretation of the data. E.J.K. and M.H. wrote the manuscript with input from all authors. M.H., E.S., and S.G. provided supervision of this study.

