## Supplemental figures and legend for "Deficiency in POLE exonuclease causes synthetic lethality in highly aneuploid cancer cells"

### Supplemental information

**Figure S1: Frequencies of each chromosomal arm loss or gain in major cancer types in the real-world database.**

**Figure S2: Associations of aneuploidy scores with additional clinical features in the real-world database.**

(A) Distribution of aneuploidy scores across tumors from patients of different genetic ancestries. AMR, Ad Mixed American; SAS, South Asian; EAS, East Asian; EUR, European; AFR, African.

(B) Distribution of aneuploidy scores across tumors with different metastasis status. Student t-tests were used for two group comparisons.

(C) Association between aneuploidy scores and patient age.

(D) Dot plots representing cooccurrence of chromosomal arm losses and gains.

**Figure S3: Association of aneuploidy scores with *TP53, AR*, *POLE* and *POLD1* genetic alterations.**

(A) Evaluation of the performance of elastic net regression performance with different alpha value. mse, mean squared error.

(B) Distribution of aneuploidy scores across tumors with different *TP53* status, separated by major cancer types in the real-world database. mut, *TP53* mutated, including short variant, structural rearrangement and homozygous deletion; wt, *TP53* wild type. Student t-tests were used for two group comparisons (mut vs. wt), ** p<0.01, *** p<0.001.

(C) Summary of sample size with each genetic alteration in the real-world database. WT, wild type; CN, copy number alteration (amplification); SV, short variant; RE, structural rearrangement.

(D) Distribution of aneuploidy scores in prostate cancer with different *AR* status in the real-world database. Student t-tests were used for two group comparisons.

(E) Distribution of aneuploidy scores across tumors with different *POLE* status in the real-world database.

(F) Distribution of aneuploidy scores across tumors with different *POLD1* status, separated by major cancer types in the real-world database. mut.exo, mutations within the POLD1 exonuclease domain (a.a.304-a.a.533); mut.other, mutations outside the POLD1 exonuclease domain; wt, wild type. Student t-tests were used for two group comparisons (mut.exo vs. wt or mut.other vs. wt), * p<0.05, ** p<0.01, *** p<0.001.

**Figure S4: Associations of aneuploidy scores with TMB, mutational signatures and microsatellite instability.**

(A) Distribution of tumor mutational burden (TMB) in tumors with different *POLE* alterations in TCGA.

(B) Distribution of TMB in tumors with different *POLD1* alterations in the real-world database.

(C) Correlation of aneuploidy scores and the mean TMB in all tumors in the real-world database (blue line: a linear regression model fit by lm() function in R)

(D) Distribution of aneuploidy scores across tumors with different mutational signatures in the real-world database.

(E) Distribution of TMB across tumors with different mutational signatures in the real-world database.

(F) Distribution of aneuploidy scores across tumors with different microsatellite instability status in the real world database. Dotted line, median aneuploidy score for POLE exonuclease mutated tumors.

(G) Distribution of TMB across tumors with different microsatellite instability status in the real world database. Dotted line, median TMB for POLE exonuclease mutated tumors.

**Figure S5: The all-in-one inducible adenine base editor.**

(A) Western blots validating the inducible expression of Cas9n(D10A)-TadA-8e(V106W) in 293, LS180 and SW837 cells.

(B) Log2 fold change of individual gRNAs tiling the POLE exonuclease domain in different cell lines.

**Figure S6: Structure of POLE highlighting the exonuclease domain and hotspot mutations P286 and V411**

(A) POLE captured on mismatched DNA, mismatch excision state (PDB 9F6L). Exonuclease domain is colored in green, catalytic residues D275 and E277 are marked in blue and represented as sticks. Template DNA strand is in plum and nascent DNA strand in orange.

(B) POLE captured on mismatched DNA, frayed substrate state (PDB 9F6K). Coloring scheme as in (A), additionally the wedge helix is colored in salmon and V411 within the wedge helix in red. Note how the wedge helix physically separates template and nascent DNA strands.

**Table S1: Excel file summarizing full results from the elastic net regression analysis**

**Table S2: Excel file summarizing tri-nucleotide motif counts on human genome and essential genes**

**Table S3: Excel file summarizing gRNA counts from the adenine base editor screen.**

### Supplemental figure 1

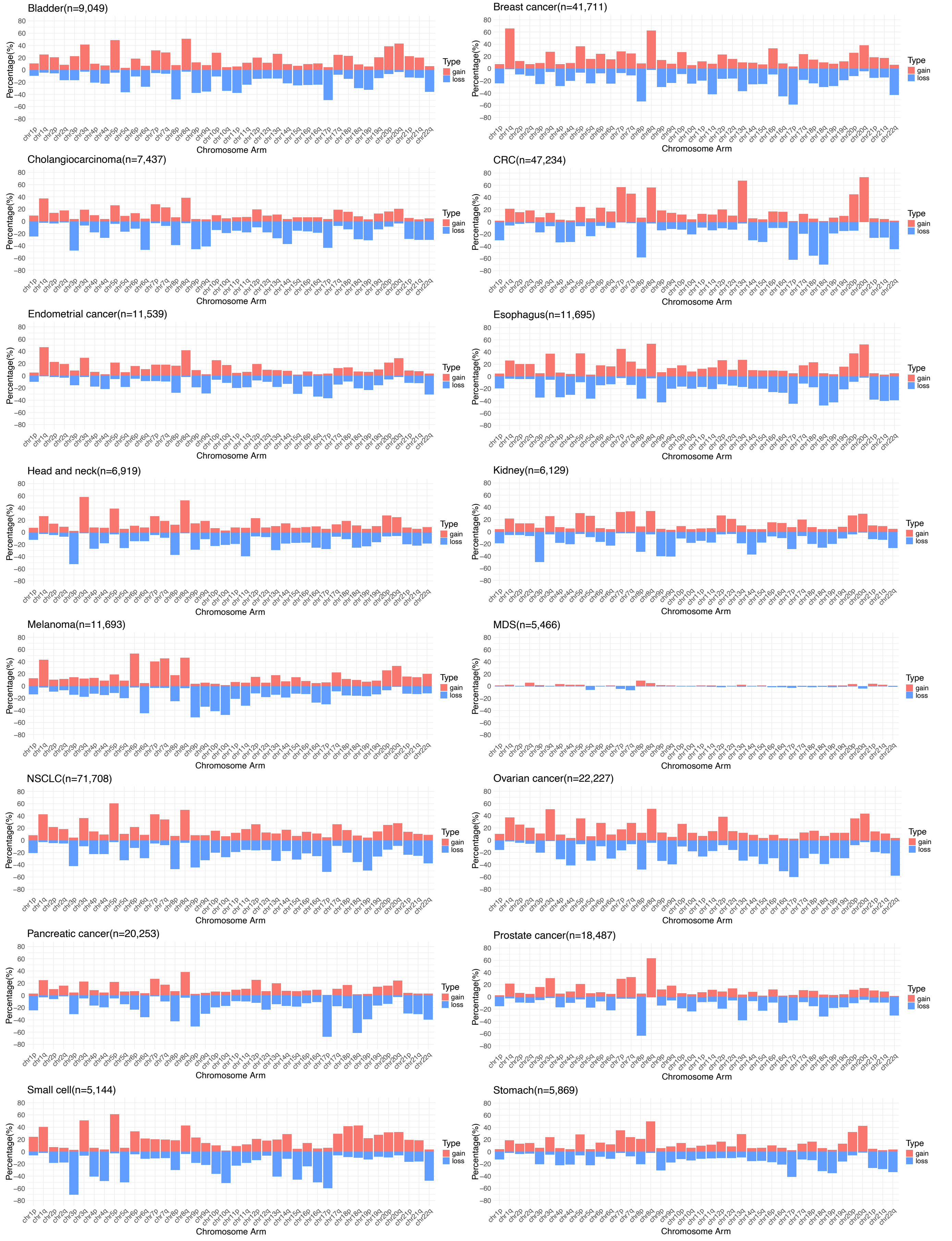

### Supplemental figure 2

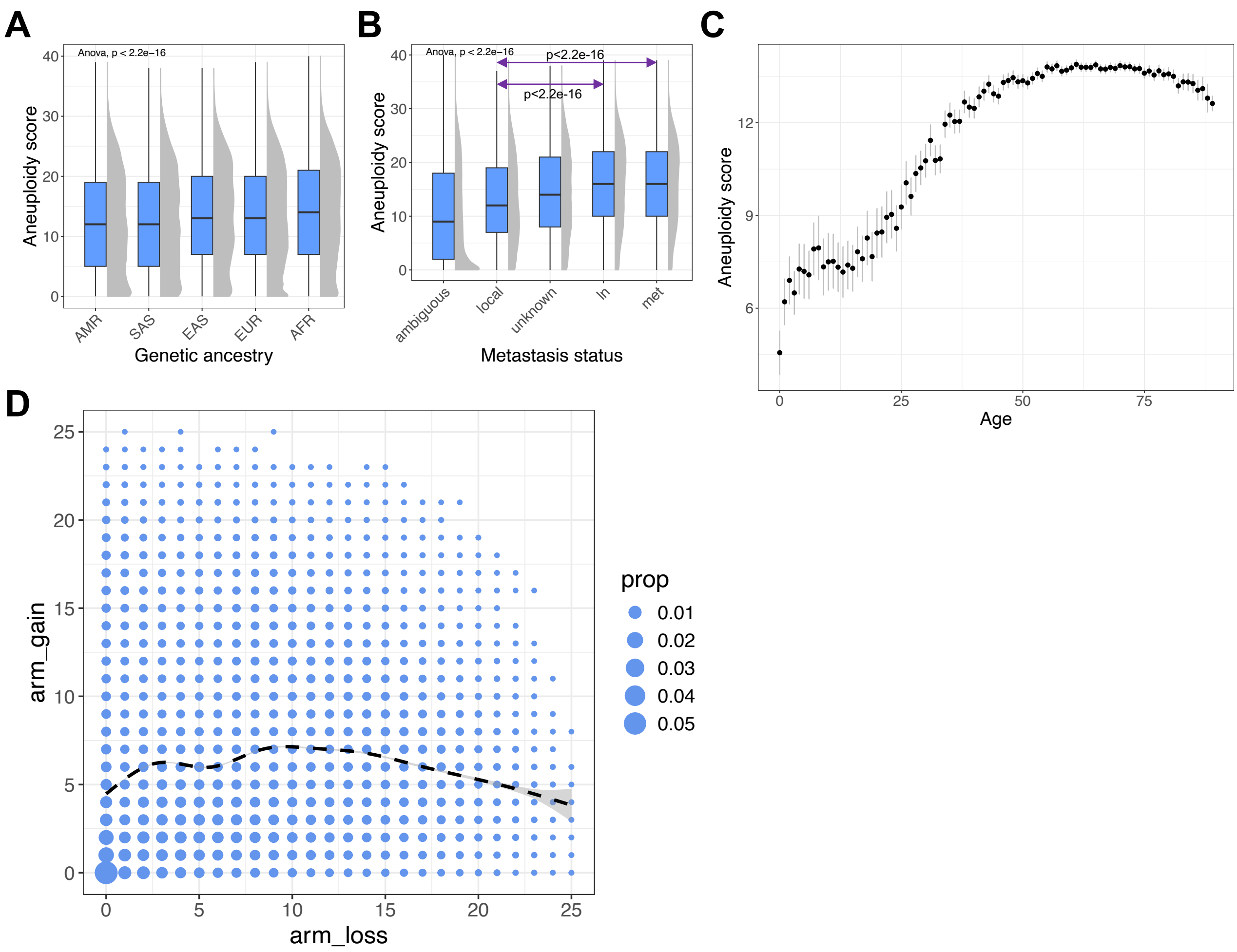

### Supplemental figure 3

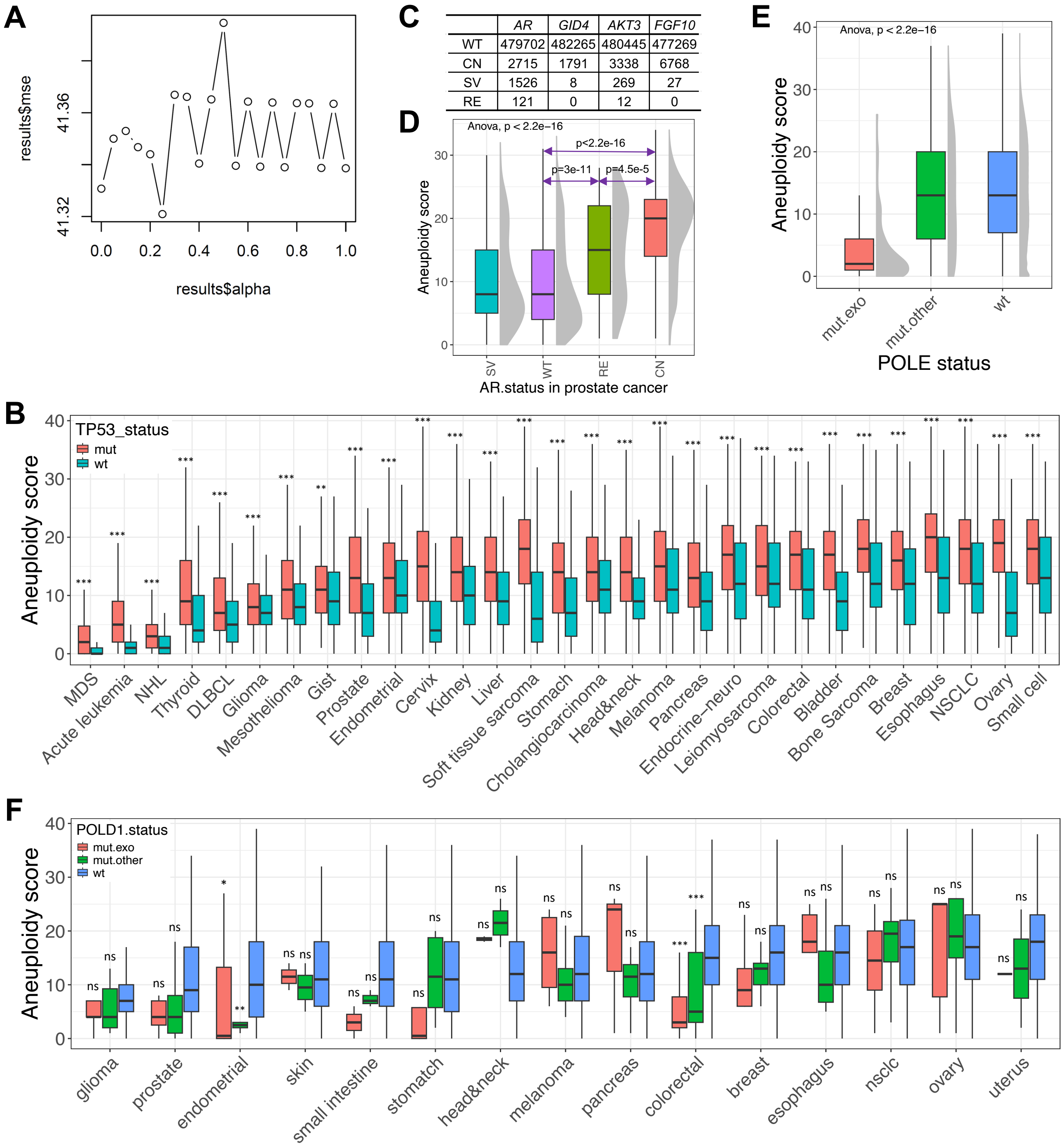


| | *AR* | *GID4* | *AKT3* | *FGF10* |
|---|---|---|---|---|
| WT | 479702 | 482265 | 480445 | 477269 |
| CN | 2715 | 1791 | 3338 | 6768 |
| SV | 1526 | 8 | 269 | 27 |
| RE | 121 | 0 | 12 | 0 |

### Supplemental figure 4

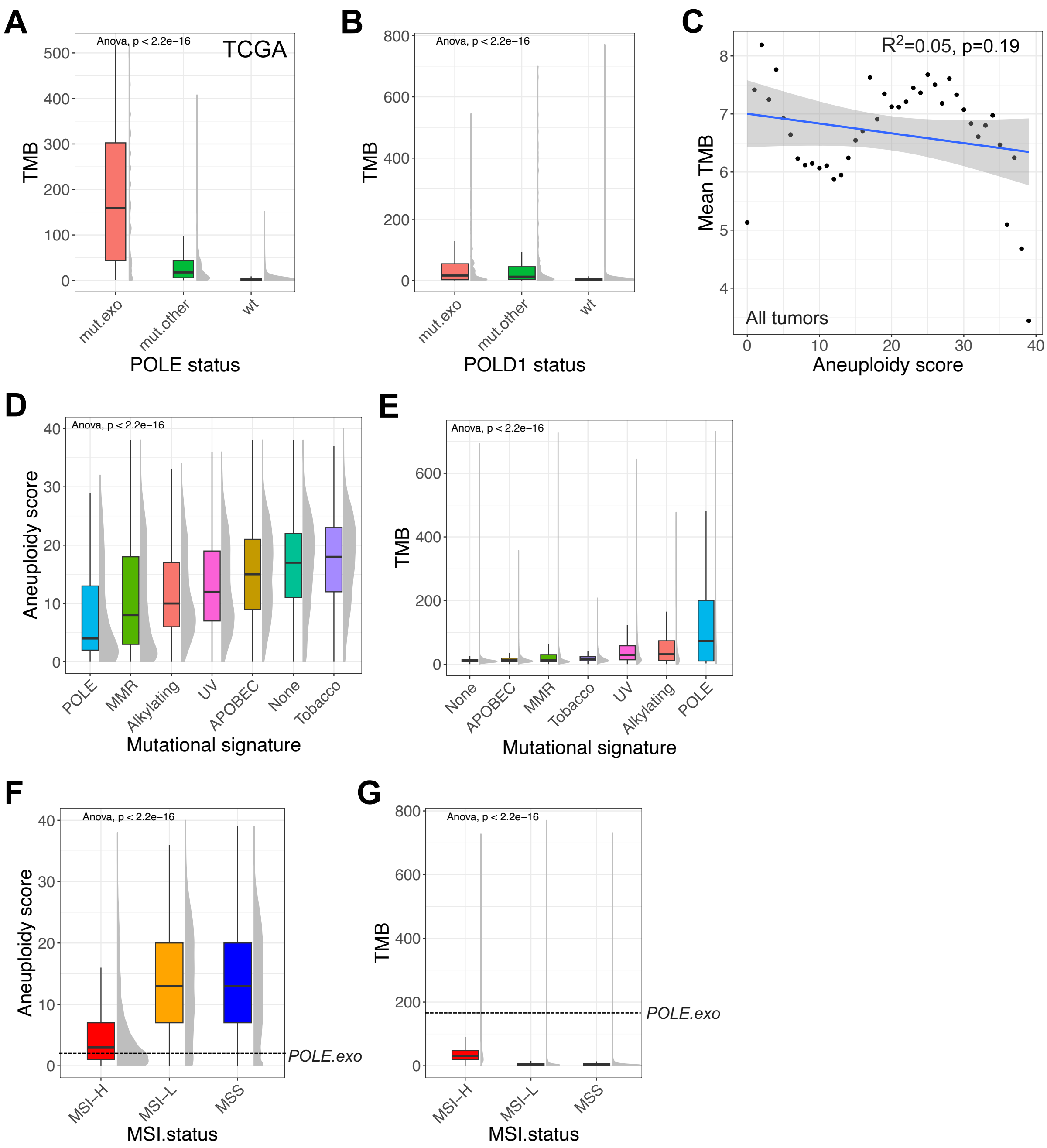

### Supplemental figure 5

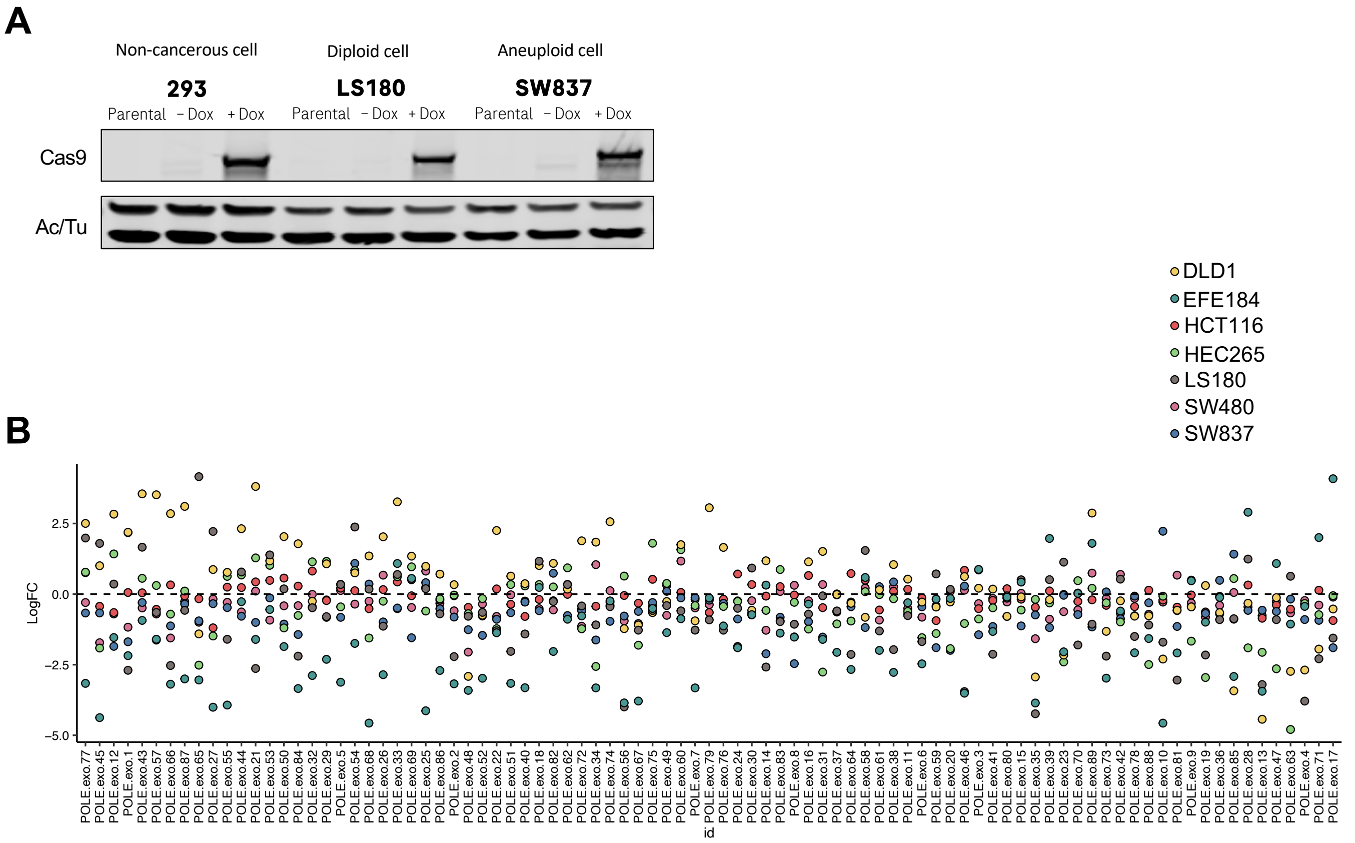

### Supplemental figure 6

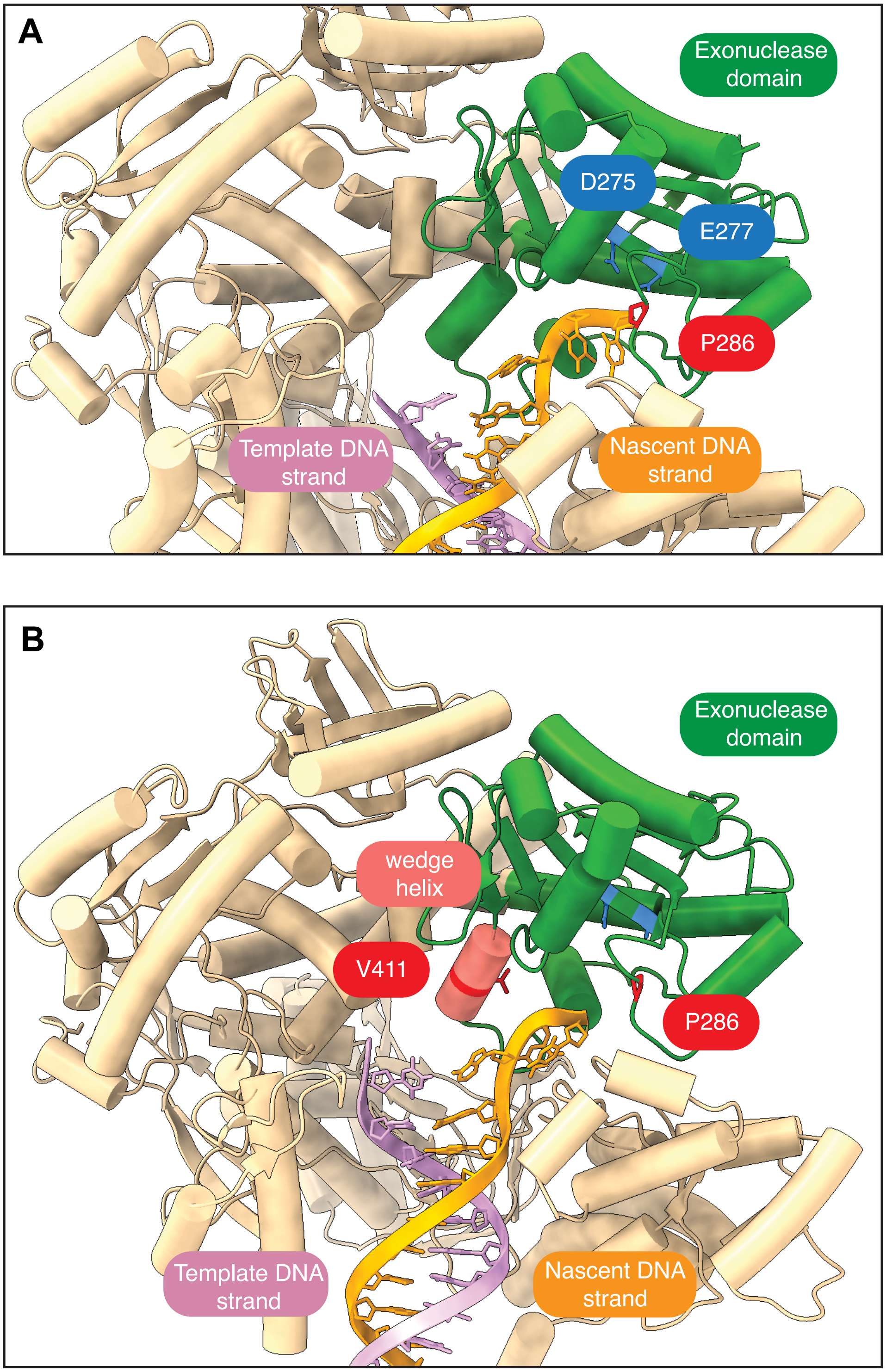
